# Cheating on cheaters dramatically affects social interactions in *Pseudomonas aeruginosa*

**DOI:** 10.1101/118240

**Authors:** Özhan Özkaya, Roberto Balbontín, Isabel Gordo, Karina B. Xavier

**Author notes:** Address correspondence to Karina B. Xavier.

## Abstract

Bacterial cooperation can be disrupted by non-producers, which can profit from public goods without paying their production cost. A cheater can increase in frequency, exhausting the public good and causing a population collapse. Here we investigate how interactions among two cheaters for distinct social traits influence the short and long-term dynamics of polymorphic populations. Using as a model *Pseudomonas aeruginosa* and its extensively studied social traits, production of the siderophore pyoverdine and the quorum sensing regulated elastase, we analyzed the social dynamics of polymorphic populations under conditions where the two traits are required for optimal growth. We show that cheaters for either trait compete with both the wild type and each other, and that mutants for pyoverdine production can prevent a drastic population collapse caused by quorum sensing cheaters. A simple mathematical model suggests that the observed social dynamics are determined by the ratio of the costs of each social trait, such that the mutant which avoids paying the highest cost dominates the population. Finally, we demonstrate how quorum sensing regulation can avoid the full loss of cooperation.

## Introduction

Bacteria are unicellular organisms, but can engage in diverse group behaviors, including biofilm formation, swarming motility, and production of extracellular proteases or iron-chelating siderophores [1–4]. The production of compounds that can benefit both producers and non-producers (public goods) can be considered as one of these cooperative behaviors. Cooperation is frequently under the threat of exploitation by cheaters: individuals that benefit from the cooperative action but contribute little or nothing to the production of the public goods. This situation, where both cooperators and cheaters can access a resource produced by the formers, is referred to as public goods dilemma [5,6]. If cheaters emerge, by mutation or migration, they can increase in frequency and cause loss of cooperation. As they rise to dominance, the public goods get exhausted and a population collapse, characterized by a strong decrease in the growth yield of the entire population, can occur; this population collapse is also referred in sociomicrobiology as ‘the tragedy of the commons’ [7–11].

Although often theorized [5,8], population collapse due to cheater expansion is hard to observe in natural populations even under conditions where cheaters spreading has been observed [12]. This raises the question of how invasion by cheaters is prevented and cooperative behaviors are maintained in microbial populations in nature. Mechanisms such as spatial structure and diffusion [13–22], pleiotropy [10,23–30], restricted migration [31], social and non-social adaptations [11,32–34], policing mechanisms [9], molecular properties of public goods [35], and metabolic strategies [36], have been proposed to play significant roles in maintaining cooperation by preventing cheater invasions and avoiding population collapse [2]. However, cheating behavior is observed *in vitro* [9–11,25], *in vivo* [37,38], and in natural populations, including clinically relevant environments such as the lungs of cystic fibrosis (CF) patients chronically infected with *Pseudomonas aeruginosa* [12,39–45].

Importantly, cheater invasion leading to the loss of the cooperative trait and the collapse of the population have been observed in laboratory studies focusing on a single trait [9–11]. However, in environments where more than one social trait is required, the roles among mutants for these traits are likely to be more complex, since a cheater for one trait can be a cooperator for another, making ‘cheater’ and ‘cooperator’ relative terms [46,47]. We hypothesize that, in environments requiring bacteria to express multiple cooperative traits simultaneously, competition among mutants for these traits can influence their social interactions and, therefore, dictate the fate of the population. To test this hypothesis, we examine the consequences of ecological interactions among two social cheaters (mutants for two different traits) and the full cooperator (the wild type) in *P. aeruginosa* populations under conditions where the two cooperative traits are required for optimal growth.

Both *lasR* and *pvdS* mutants have been studied individually in a large number of sociomicrobiology studies [10,25,35,48–52], and are among the most common mutants recurrently isolated from the sputum samples of CF patients [12,39,41].

LasR is the master regulator of quorum sensing and, among many other genes, controls the production of extracellular elastase [53–55], which is essential for *P. aeruginosa* to use complex sources of amino acids, such as casein, as a carbon and nitrogen source [56–59]. Previous studies showed that *lasR* mutants grow poorly in media containing casein as the only carbon source, but increase in frequency when mixed with wild-type (WT) bacteria. Such increase leads to a population collapse where total cell numbers are drastically reduced due to depletion of producers of the public good [9–11].

PvdS is an alternative sigma factor, that among other genes, controls the transcription of genes responsible for pyoverdine biosynthesis [60,61]. In iron-limited environments, *P. aeruginosa* can secrete pyoverdine, which binds iron from the environment and is subsequently retrieved, providing iron to the cell [49]. Mutants in pyoverdine synthesis (e.g. *pvdS*) do not pay the cost of its production but are still able to retrieve the iron-bound pyoverdine produced by others, gaining a fitness advantage and increasing in frequency in mixed populations [35,62,63].

We analyzed the cheating of a *lasR* knock-out (KO) mutant over wild-type bacteria in an environment where production of elastase is required (medium with casein as the sole carbon source with iron supplementation). Then, we modified the conditions (medium with casein as the sole carbon source with iron depletion by human apo-transferrin) to cause pyoverdine production to be also required and added a third social player (a *pvdS* KO mutant), to study the interactions in these two different scenarios in short and long-term competition experiments. We found that the fitness advantage of the *lasR* mutant disappears when the *pvdS* mutant is in the culture and the two traits are necessary. The long-term consequence of the interaction between these two mutants is the prevention of the drastic population collapse, which occurs (*i*) irrespectively of the presence of *pvdS* under conditions where only elastase is required, or (*ii*) in the absence of the *pvdS* in conditions where the two traits are required. The observed dynamics can be explained by a simple mathematical model of multiple public good competition, which predicts the dominance of the mutant that avoids expressing the trait with the highest cost, eventually causing the corresponding population collapse associated with the loss of that trait.

## Results

### Cheating capacity of lasR mutant depends on abiotic and biotic factors

We first investigated the growth yields of the WT and *lasR* and *pvdS* mutants in monocultures under environmental conditions where either trait, both, or none are required. In medium where casein is the sole carbon source, iron-supplemented casein medium (Casein + Fe), and therefore elastase is required, the *lasR* mutant has a lower growth yield than the WT and the *pvdS* mutant (Figure 1A). In medium strongly depleted in iron, as it has been repeatedly shown in the literature, the growth yields of all the three strains, including the WT, are severely reduced [34,35,63–67]. However, the growth yield of the *pvdS* mutant is significantly lower than those of the WT and the *lasR* mutant in iron depleted media with casamino acids (CAA + Transferrin), where only pyoverdine is necessary (Figure 1B). In a medium with casein as the sole carbon source and low iron, namely, iron-depleted casein medium (Casein + Transferrin), where both traits are required, both *lasR* and *pvdS* mutants have a lower growth yield than the WT (Figure 1C). In this medium, the growth yield of the *lasR* mutant is smaller than that of the *pvdS* mutant, indicating that the traits have different benefit/cost ratios in this environment. Importantly, the WT and the two mutants have similar growth yields in a medium where none of the traits are required, iron-supplemented CAA medium (CAA + Fe) (Figure 1D), in accordance with the expectation that the differences in growth yields of the mutants across media are due to the lack of expression of each social trait in the corresponding mutant. Moreover, the observation that the *lasR* mutant only has a growth yield significantly lower than the WT in the media with casein as the sole carbon source indicates that, even though LasR regulates many genes besides those responsible for elastase production [53–55], most do not significantly affect fitness under the conditions tested (For more details about the media used in this study and the growth yield differences of the strains, see Supplemental Information).

**Figure 1.**
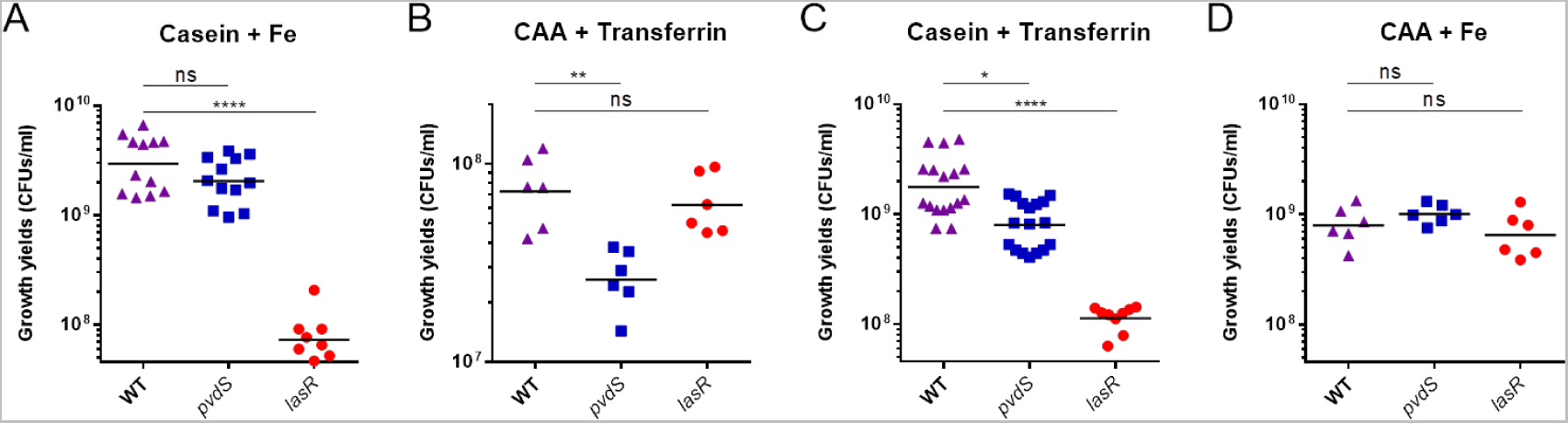
P. aeruginosa lasR and pvdS mutants have lower growth yields than WT in media where elastase and/or pyoverdine are required, respectively. Growth yields (CFU/ml) of WT (purple triangles), *pvdS* (blue squares), and *lasR* (red circles) strains of *P. aeruginosa* monocultures after 48 hours of incubation in **(A)** iron-supplemented casein medium (Casein + Fe), **(B)** iron-depleted casamino acids medium (CAA + Transferrin), **(C)** iron-depleted casein medium (Casein + Transferrin) and **(D)** iron-supplemented casamino acids medium (CAA + Fe). Each data point represents an individual biological replicate (N≥6) and the horizontal bars indicate the means of each group. For comparisons, Kruskal-Wallis test with Dunn’s correction was used, ns=not significant P>0.05, * P≤0.05, ** P≤0.01, *** P≤0.001, **** P≤0.0001.

We then determined the relative fitness of each mutant in competition with the WT and each other in the different media (Figure 2). When co-cultured, at a ratio WT*+lasR* of 9:1, in conditions where only elastase production is required, the *lasR* mutant can cheat on the WT, since *lasR* increases in frequency with respect to it (Figure 2A-left), whereas such increase does not occur when elastase production is not required (Figure 2G). The introduction of the *pvdS* mutant (at ratio WT*+lasR+pvdS* of 8:1:1) does not significantly affect the cheating behavior of the *lasR* mutant, since *lasR* can also increase in frequency in the triple co-culture (Figure 2A-right). The incapability of the *pvdS* mutant to affect cheating by *lasR* is consistent with the fact that *pvdS* does not cheat under these conditions (Figure 2B). As expected, *pvdS* can cheat in medium where only pyoverdine is required, whereas *lasR* cannot (Figure 2, panels D and C, respectively). Next, we studied the interaction between these social players in conditions where the two traits are necessary. In these conditions, the *lasR* mutant also increases in frequency in co-culture with the WT (Figure 2E-left). Remarkably, the introduction of the *pvdS* mutant under these conditions causes the cessation of cheating by *lasR* (Figure 2E-right), which is consistent with the fact that *pvdS* can cheat on the WT in co-culture (Figure 2F-left) and both on the WT and the *lasR* mutant in triple co-culture (Figure 2F-right, Figure S1F). Importantly, in conditions where neither of the social traits are necessary, no cheating can be observed (Figure 2G-H), further ratifying that the effects observed are due to social interactions.

**Figure 2.**
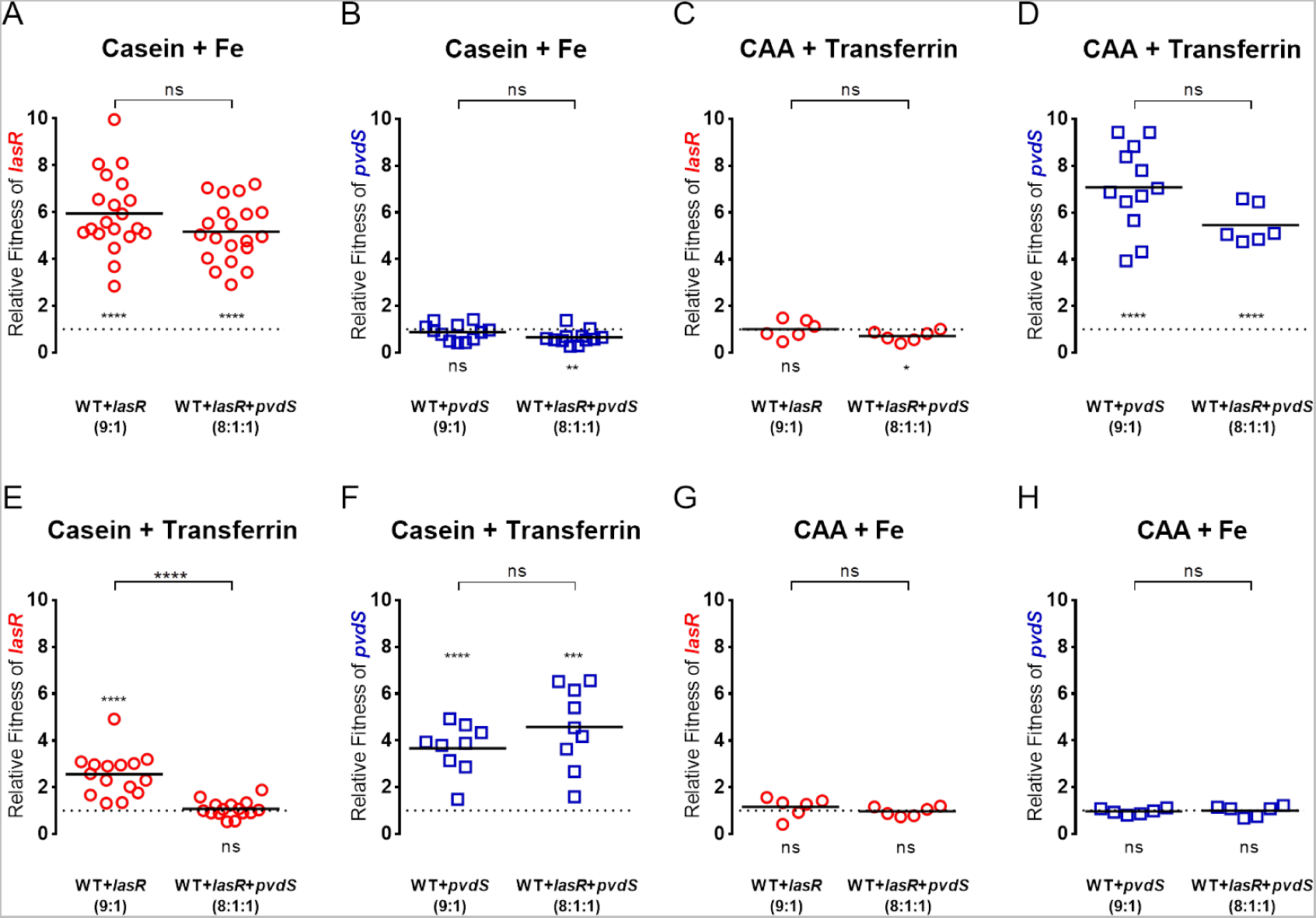
Relative fitness of lasR or pvdS in various media in double or triple co-cultures. Relative fitness *(v)* of *lasR* (red circles in **A, C, E**, and **G**) or *pvdS* (blue squares in **B, D, F,** and **H**) were calculated as the change in frequency of each mutant relative to the rest of the strains in each culture after 48 hours of incubation in **(A** and **B)** iron-supplemented casein medium (Casein + Fe), **(C** and **D)** iron-depleted casamino acids medium (CAA + Transferrin), **(E** and **F)** iron-depleted casein medium (Casein + Transferrin), and **(G** and **H)** iron-supplemented casamino acids medium (CAA + Fe). Relative fitness of *lasR* (**A, C, E,** and **G**) was calculated in co-cultures with WT, or with WT and *pvdS*. Relative fitness of *pvdS* (**B, D, F,** and **H**) was calculated in co-cultures with WT, or with WT and *lasR*. Initial ratios of the strains in each co-culture are 9:1 for WT+*lasR* and WT+*pvdS*, and 8:1:1 for WT+*pvdS*+*lasR*. Mann-Whitney two-tailed test was used to compare the relative fitness values of each mutant in double and triple co-cultures (significance symbols are located at the middle-top of each plot above the brackets). Dotted lines indicate *v*=1. Relative fitness values above the dotted lines (*v*>1) indicate that the strain is cheating and below the dotted lines (*v*<1) indicate that the strain is being cheated. One-sample t-test was used to determine whether each dataset is significantly different than 1 (significance symbols are located above the dotted line when *v*>1 and below the dotted line when *v*≤1). Each data point indicates an individual biological replicate (N≥6) and horizontal lines indicate the means of each group. ns=not significant P>0.05, * P≤0.05, ** P≤0.01, *** P≤0.001, **** P≤0.0001.

Notably, *lasR* mutants have been reported to produce lower amounts of pyoverdine than the WT in iron-limited succinate minimal medium [68]. However, we found no significant difference in pyoverdine production between WT and *lasR* across the different media used in this study (Figure S2). The difference between our results and those of Stintzi and colleagues might be due to differences in the media used in the two studies [64], or potential differences in the strains used.

Altogether, these results demonstrate that the cheating capacities of the two social mutants studied here are context-dependent, varying not only with the environment, but also with the level of polymorphism in the population.

### Invasion of lasR mutant leads to a drastic collapse of the population

We next asked what are the long-term consequences of the different cheating capacities of *lasR* for the overall fitness of the population by performing long-term propagations (Figure 3). We started co-cultures of WT+*lasR* or WT+*lasR*+*pvdS* (at 9:1 and 8:1:1 initial ratios, respectively), either in medium requiring only elastase production (Figure 3A and B), or in medium where the two traits are needed (Figure 3C and D). Propagations were performed by transferring an aliquot of each culture to fresh media every 48 hours. Before each passage, growth yields and frequencies of WT, *pvdS*, and *lasR* cells were determined (Figure 3).

**Figure 3.**
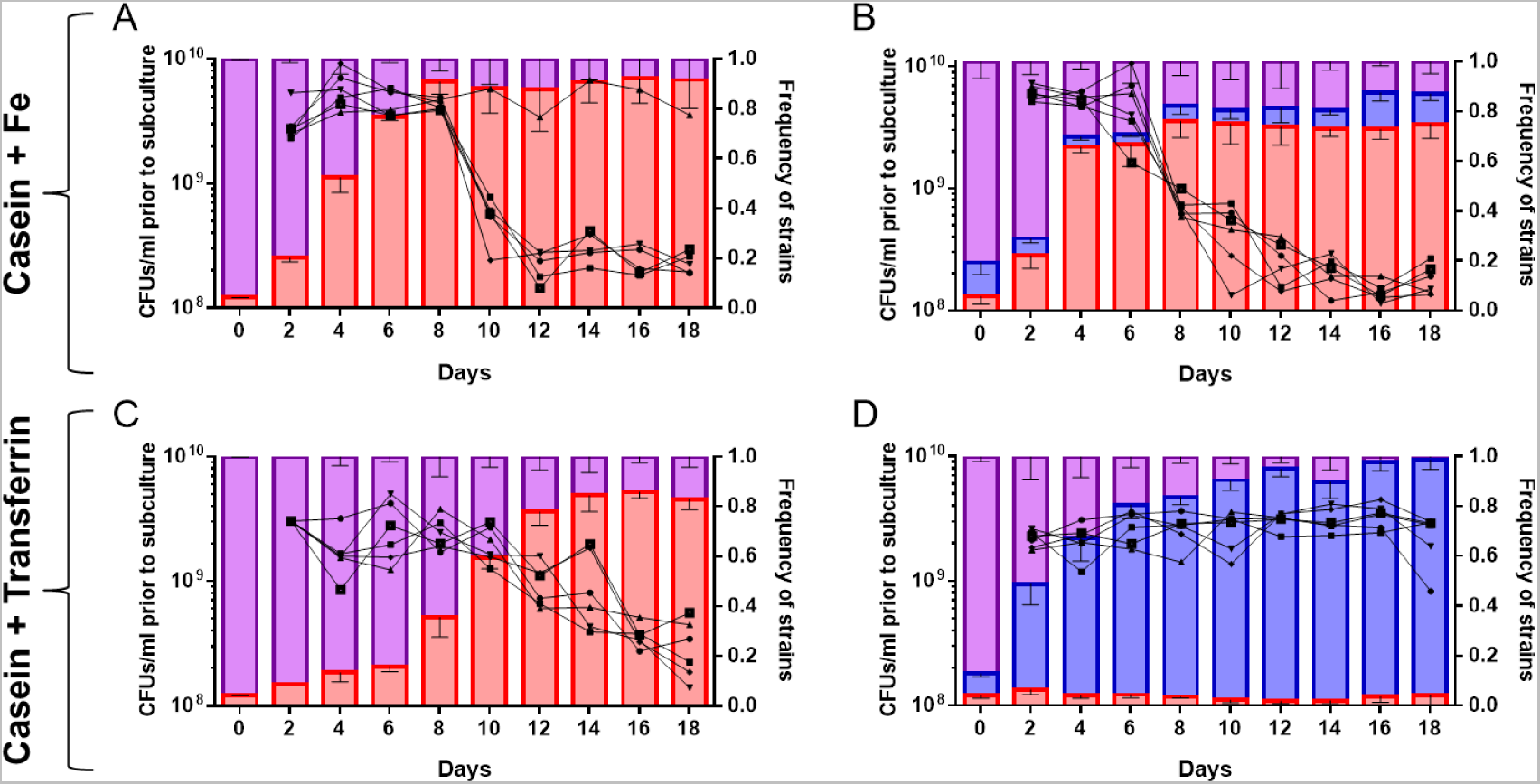
Effects of abiotic and biotic factors on growth yields and strain composition of the population in long-term propagations. Left Y-axes represent CFU/ml values prior to subculture; black symbols correspond to the CFU/ml values of each of the 6 biological replicates tested for each condition. Right Y-axes show the frequencies of WT (purple), *lasR* (red), and *pvdS* (blue) at each time point; data are shown as bars and represent the means of 6 biological replicates, error bars indicate SD. X-axes show the days of propagations. **(A)** WT and *lasR* co-cultures mixed at an initial frequency of 9:1 in iron-supplemented casein media. **(B)** WT, *lasR*, and *pvdS* triple co-cultures mixed at initial an initial frequency of 8:1:1 in iron-supplemented casein media. **(C)** and **(D)** same as in (A) and (B) but in iron-depleted casein media.

We observed that, in five out of six replicates of WT*+lasR* co-cultures in the medium where only elastase is required, the *lasR* mutant quickly increased in frequency throughout the first 8 days (4 passages), reaching up to 90% of the population (red bars in Figure 3A). The total cell numbers of the populations (black lines) rapidly decreased by day 12, and no recovery was observed in subsequent passages (Figure 3A). We defined this drastic decrease in density as the population collapse caused by the domination of the *lasR* mutant. One replicate, out of six, did not follow this trend; in this case, no population collapse was observed, and the total cell numbers remained high throughout the experiment (Figure 3A). The fact that this only occurred in one of the six replicates suggests that, in this particular replicate, the WT may have acquired non-social beneficial mutation(s) that could prevent invasion of the *lasR* mutant, as it was described in a recent study [11].

Next, we analyzed long-term competitions in triple co-cultures (WT, *pvdS*, and *lasR*; with initial frequencies of 8:1:1, respectively) in the medium where only elastase is required (Figure 3B). In this case, we observed an increase in *lasR* frequency similar to that of seen in WT+*lasR* co-cultures (Figure 3A), which was also accompanied by a drastic decrease in the overall population size. At day 12 of the propagation, all 6 populations collapsed (Figure 3B). The frequencies of the *pvdS* mutant varied between 4% and 15% throughout the experiment, with no indication of any sustained increase (blue bars in Figure 3B). This result is consistent with the predictions from the relative fitness measurements, which shows no cheating of *pvdS* in these conditions (Figure 2B).

Then we propagated WT+*lasR* co-cultures in the medium where the two traits are necessary (Figure 3C). In these propagations, the *lasR* mutant also increases in frequency throughout the first days, but at a slower pace than when only elastase is required (compare panels A and C in Figure 3). The total cell numbers remain high until days 10-12, but, as the *lasR* frequencies increase to about 80%, the density of the population decreases, collapsing by day 18. Hence, in all the three scenarios described above, the dominance of the *lasR* mutant, which presumably resulted in the exhaustion of the public good elastase, caused a drastic population collapse (Figure 3A–C).

### pvdS prevents the drastic population collapse caused by the invasion of the lasR mutant

Our short-term competitions revealed that the cheating capacity of *lasR* is influenced not only by abiotic, but also by biotic conditions, as the presence of *pvdS* under conditions where both traits are required reduces the relative fitness of the *lasR* mutant (Figure 2C). Therefore, we investigated if, in propagations in the medium where both traits are needed, *pvdS* could prevent the drastic population collapse caused by *lasR* invasion. Indeed, Figure 3D shows that *lasR* does not increase in frequency, staying at approximately 3% throughout the experiment. In contrast, *pvdS* rapidly expands to an average frequency of 96% at day 18. As the *lasR* mutant does not increase in frequency, cell densities of the overall populations do not decrease and collapse of the population is not observed.

Given that *pvdS* dominated the populations, a reduction in cell numbers due to its invasion could be expected. Indeed, full fixation of the *pvdS* mutant should result in a small population decrease close to the levels of the *pvdS* mono-cultures (Figure 1C). However, at the end of the propagation experiments (day 18), complete fixation of *pvdS* had not yet been reached, and an average of 4% of WT and *lasR* cells were detected in the populations (Figure 3D). We hypothesized that the presence of only 4% of pyoverdine producers in the population could be enough to sustain the growth of the entire populations to levels similar to the WT mono-cultures similarly to what has been reported recently for cultures in chemostat [69]. The results shown in Figure S3A support this hypothesis, since the growth yields of WT+*pvdS* mixed cultures at different initial frequencies of *pvdS* significantly decrease only when the initial frequency of *pvdS* is 98% or less, whereas mixtures with 3-4% of WT cells (or WT and *lasR* cells) have growth yields similar to that of WT monocultures. These results demonstrate that a small proportion of pyoverdine producer cells (WT and/or *lasR* cells) are sufficient to produce enough pyoverdine to sustain the entire population. This justifies why, in the propagation shown in Figure 3D, where at day 18, *pvdS* reached an average frequency of 96%, no significant drop in cell numbers was observed. Moreover, these results indicate that, if the propagations were to continue, *pvdS* fixation and the subsequent decrease in cell density could be expected. In fact, new WT+*pvdS* and WT*+pvdS+lasR* propagations with much higher initial frequencies of *pvdS* (75-85%), allowed to observe this population collapse (Figure S3B and C). The reason why *lasR* or *pvdS* domination lead to a stronger or milder population collapse, respectively, is related with the different characteristics of these two mutants, which have different fitness in mono-culture in the medium requiring both traits (Figure 1C), presumably as a consequence of the differences in cost and benefits of the traits involved.

Remarkably, the presence of the *pvdS* mutant in the 3-way competition under conditions where the two traits are required has a strong effect on the outcome of the propagations in terms of growth yields, which is dramatically different from those of the other three scenarios tested, since *pvdS* domination prevents the drastic population collapse (Figure 3D) caused by the expansion of *lasR* in the other three conditions (Figure 3A–C). This occurs because, in this environment, although the *lasR* mutant is still being able to cheat on the WT (Figure S1C), it is being cheated by *pvdS* (Figure S1D).

Importantly, long-term propagation experiments of WT*+lasR* and WT*+lasR+pvdS* in medium where neither of the traits are required showed no significant change in the population densities (Figure S4).

### Alterations in carbon or iron source availability can prevent or induce the collapse

We reasoned that, if social interactions dominate over *de novo* adaptive mutations in long-term dynamics, alterations of the abiotic factors in the triple cultures should modify the social role of each mutant (by changing the costs and benefits of the cooperative traits) and therefore affect the outcome for the populations. Indeed, in the propagation of WT*+lasR* co-cultures in the medium where only elastase is required, changing the carbon source from casein to CAA during the course of the propagation (making elastase unnecessary) eliminates the advantage of the *lasR* mutant, and this environmental change is sufficient to prevent the population collapse (Figure 4A). Conversely, addition of iron to the medium where both traits are required (thus making pyoverdine unneeded) reverts the expansion of the *pvdS* mutant, favoring *lasR* cheating, and ultimately causing the collapse of the populations at day 18 (Figure 4B). We confirmed that changes in final frequencies observed in Figure 4B were not due to the high initial frequencies of *pvdS*, because even though the selective advantage of *pvdS* is frequency dependent, this mutant is capable of cheating even at frequencies higher than 90% (Figure S5).

**Figure 4.**
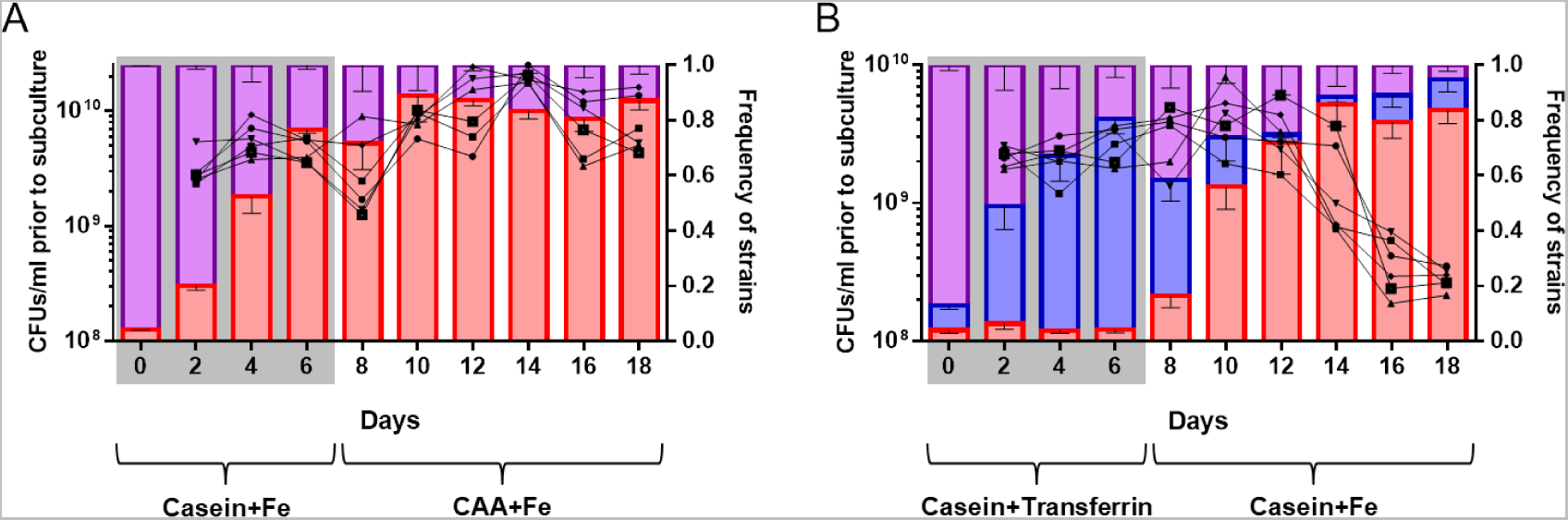
Effects of alterations of abiotic conditions in long-term propagations. **(A)** After the 6^th^ day of the competitions of WT+*lasR* co-cultures in iron-supplemented casein medium (Casein + Fe) (Figure 3A), aliquots were transferred into iron-supplemented CAA medium (CAA + Fe) to relieve the requirement for digesting casein by elastase production (N=6, data from the first 6 days are from Figure 3A). **(B)** After the 6^th^ day of the competitions of WT+*lasR*+*pvdS* triple co-cultures in iron-depleted casein medium (Casein + Transferrin) (Figure 3D), aliquots were transferred into iron-supplemented casein medium (Casein + Fe) to relieve the requirement for pyoverdine production (N=6, data from the first 6 days are from Figure 3D). Legends as in Figure 3.

Overall, these results confirm the cheating role of the two mutants, and also demonstrate the preponderance of social interactions over evolutionary adaptation by *de novo* mutation in the propagation experiments shown in Figure 3.

### A mathematical model of a 3-way public goods game explains the dynamics of the cheating mutants

To further investigate the general factors determining the dynamics of competitions among cooperators and cheaters we built a simple mathematical model (see Supplemental Information, Mathematical Model 1 and 2). The model assumes that the cost (*c*) of a cooperative trait is lower than the benefit (*b*) associated with this trait (*b*>*c*>0), and also that the benefit provided by the cooperative trait is equal for the entire population, as it would be expected in the case of an equally accessible public good in a well-mixed environment. Spatial structure, diffusion, or privatization, which would alter the benefit gained from the public good for cooperators and cheaters asymmetrically, were not considered in the model. The parameters used are described in Supplementary Table S1. As can be seen from the fitness definitions of the three players involved in our simple 3-way public model (Supplemental Information, Mathematical Model 1 (equations 1 to 3)), the cheaters always have a higher fitness than the cooperator due to the costs (*c_1_* or *c_2_*) saved. Figure 5A shows the predicted mean fitness (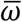) and final frequencies of the different strains in the population assuming different *c*_*1*_/*c*_*2*_ ratios. It can be easily seen that cooperators will always go extinct, and that the two cheaters can only co-exist when *c_1_* = *c_2_*. Whenever *c_1_* ≠ *c*_*2*_, then the cheater that produces the trait with highest cost will lose. Therefore, the relation between *c*_*1*_ and *c*_*2*_ determines which cheater will dominate the population, independently of the benefits (*b*_*1*_ and *b*_*2*_) of these cooperative traits. On the other hand, the mean fitness, 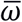, is affected by the difference between *b* and *c* values of each trait.

**Figure 5.**
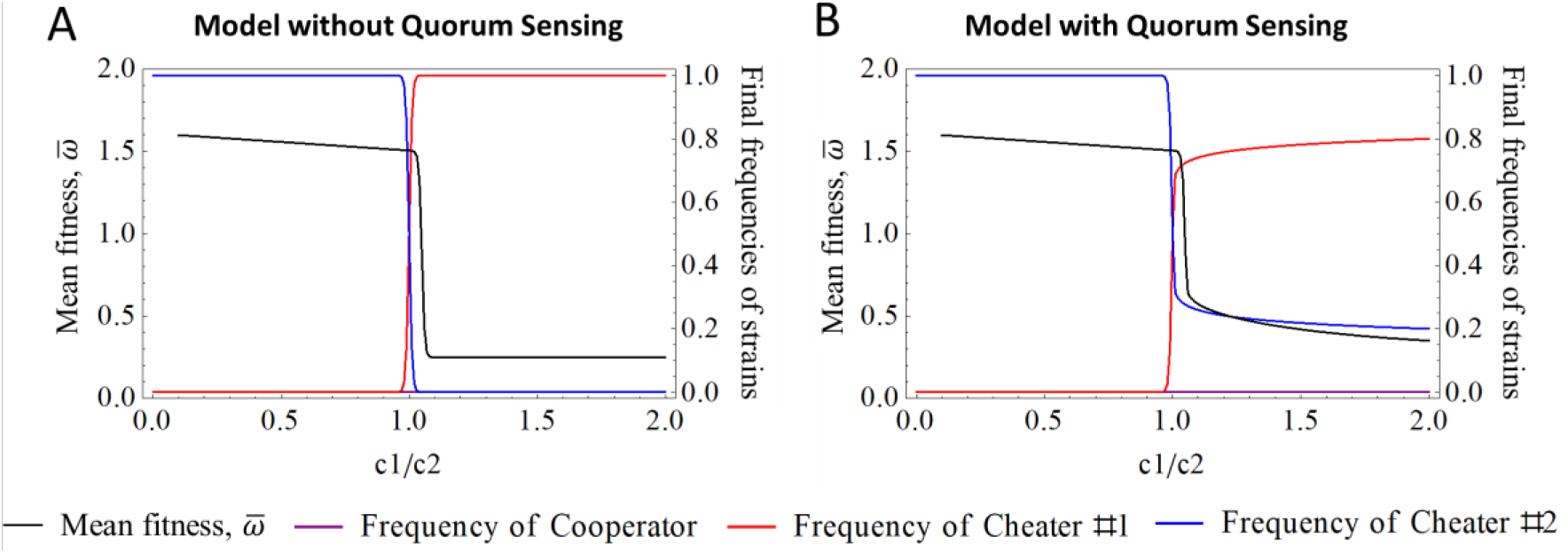
Mathematical model for the final frequencies of the three strains in relation to the ratio of *c*_*1*_/*c*_*2*_. In Left-Y axes, the mean fitness, 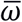, is shown in black. In Right-Y axes, frequencies of cheater of the 1^st^ cooperative trait (red), cheater of the 2nd cooperative trait (blue), and cooperator of both cooperative traits (purple) are shown in relation to the ratio of *c*_*1*_/*c*_*2*_ (X-axes) either without **(A) - mathematical model 1** or with the influence of quorum sensing (QS) regulation on the 1^st^ cooperative trait **(B) - mathematical model 2**. The values given to the parameters of the simulations are: *p*_*coop*_(0)=0.8, *p*_*ch1*_(0)=0.1, *p*_*ch2*_(0)=0.1, 0.001≤*c*_*1*_<0.199, *b*_*1*_=1.5, *c*_*2*_=0.1, *b*_*2*_=0.25, ω_0_=0.1, time (as arbitrary units of cumulative numbers of cell divisions)=1800. In (B) the 1^st^ cooperative trait is regulated by QS with *n*=30 and *th*=0.8.

We simulated the four scenarios corresponding to the conditions in Figure 3. As shown in Figure S6 in panels A and C, the cooperator for both traits and the cheater of the 1^st^ cooperative trait compete, while the cheater of the 2^nd^ cooperative trait is absent, whereas in panels B and D all three strains compete. In panels A and B, only the 1^st^ cooperative trait is necessary, while in panels C and D both traits are required. The *c*_*1*_/*c*_*2*_ ratios defined in the model are estimated from the ratios of the relative fitnesses determined in the competitions shown in Figure 2E and F. The results of the model for the four scenarios resemble the experimental data, explaining changes in frequencies reasonably well (Figure S6A-D). However, this simple model predicts complete fixation of the expanding mutant (Figure S6A-D), and cannot explain the lack of fixation of *lasR* (Figure 3A-C) or *pvdS* (Figure 3D) observed experimentally. As discussed above, as long as *de novo* mutations are not acquired, *pvdS* can reach fixation when co-cultured either with WT, or with WT and *lasR* under conditions where the two traits are needed (Figure S3B and C), which is in accordance to the model. However, that was not the case when *lasR* expansion was observed. We tested experimentally whether fixation of the *lasR* mutant could occur if the propagations were continued under conditions where *lasR* was expanding. Our results show that, when we initiate WT+*lasR* competitions at initial *lasR* frequencies similar to those at day 18 in Figure 3A, *lasR* still fails to reach fixation (Figure 6A). This dynamical behavior of *lasR* is not predicted under the assumptions of the model and suggests that other processes are taking place in the experiment.

**Figure 6.**
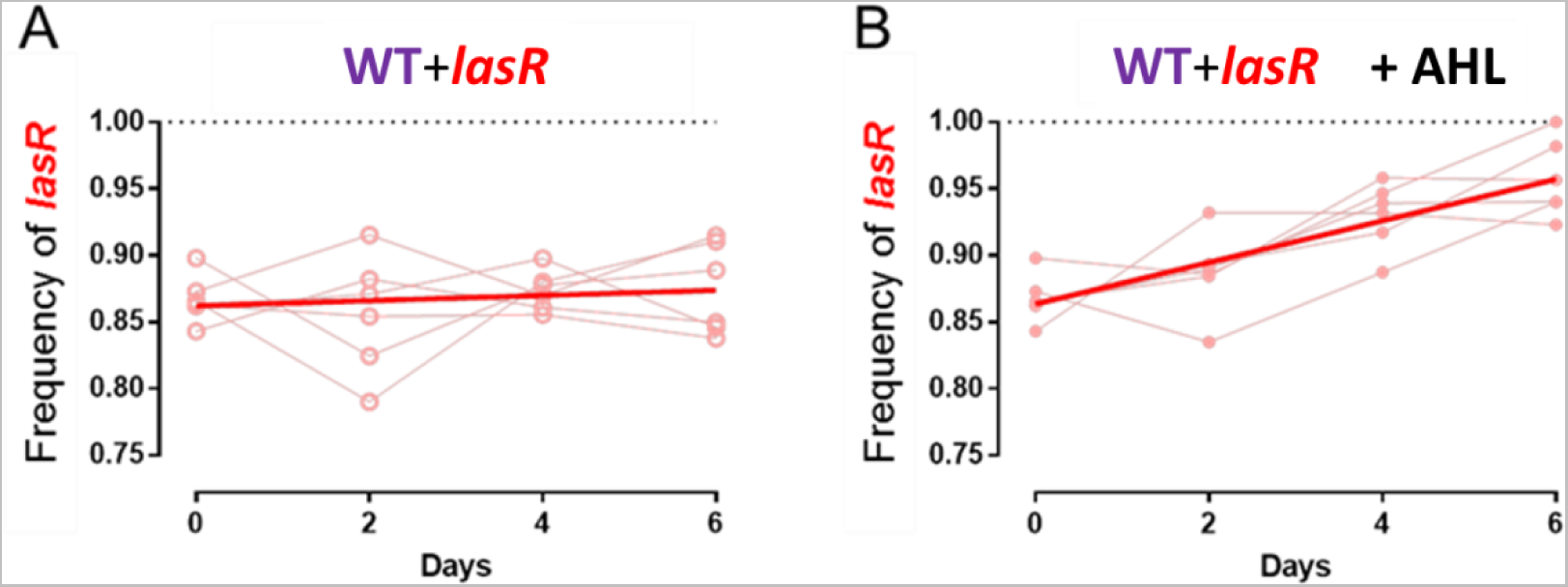
Frequencies of lasR in propagations of WT+lasR co-cultures in iron-supplemented casein media in the absence or presence of exogenously added quorum sensing signal AHL. Initial frequencies of *lasR* of 80-90% were used, these frequencies were similar to those of the 18^th^ day in Figure 3A. Cultures were propagated throughout 6 days by passing the fresh media each 48 hours. **(A)** Frequency changes of *lasR* in WT+*lasR* co-cultures (red). **(B)** is the same as (A) but with 5μM AHL (3OC_12_-HSL) added to the media. Red lines indicate linear regressions. Dotted lines represent 100% domination of *lasR*.

Given that the *lasR* gene and elastase production are regulated by quorum sensing, we hypothesized that quorum sensing could be responsible for the lack of fixation of *lasR* mutant observed experimentally. Quorum sensing regulation should reduce both the cost and the benefit of elastase production when the cooperators are below the quorum sensing threshold, as cells will not produce elastase in that phase. We therefore modelled the effect of quorum sensing on fitness equations by assuming a Hill function where the cost and benefit of the 1^st^ cooperative trait are sharply reduced when the frequency of the cheater for the 1^st^ cooperative trait reaches a given threshold value (see Supplemental Information, Mathematical Model 2 for the model including quorum sensing). In this case, fixation of the mutant for the 1^st^ cooperative trait can only happen if *c*_*1*_<*c*_*2*_. When *c*_*1*_≥*c*_*2*_, both cheaters can co-exist in the population (Figure 5B). As shown in Figure 7, the simulations of the modified model including quorum sensing for the four experimental conditions predict accurately their frequency dynamics. Moreover, it also predicts that mutants for traits regulated by quorum sensing, like *lasR*, will not reach fixation.

**Figure 7.**
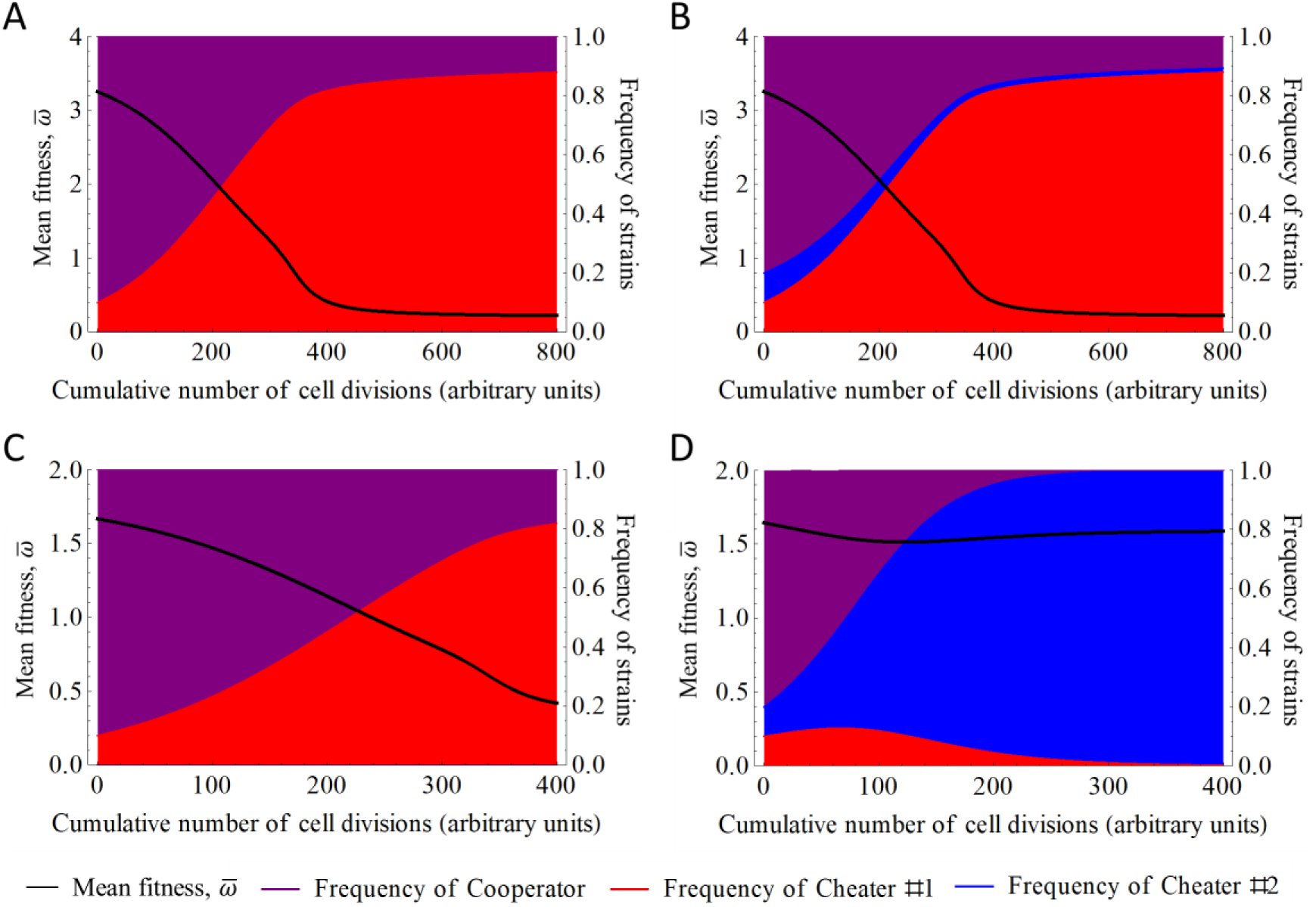
Results of the mathematical model simulating the four scenarios in Figure 3. Model includes quorum sensing regulation of the 1^st^ cooperative trait (*b*_*1*_ and *c*_*1*_ are negatively regulated via a Hill equation as a function of the frequency of the mutant of this trait, *p*_*ch1*_). Left Y-axes show 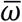, the mean fitness of the entire population which is a function of *b* and *c* values; these values correspond to the biomass gain due to benefiting from the cooperative action (*b*), and the energy spent to the cooperative action instead of biomass increase (*c*). Right Y-axes show the frequencies of *p*_*coop*_ (e.g. WT, purple), *p*_*ch1*_ (e.g. *lasR*, red) and *p*_*ch2*_ (e.g. *pvdS*, blue). X-axes show the cumulative numbers of cell divisions as arbitrary units. In panels **(A)** and **(B)**, only the 1^st^ cooperative trait is necessary (*b*_*1*_ > *c*_*1*_ > 0, whereas *b*_*2*_ = *c*_*2*_ = 0), while in panels **(C)** and **(D)** both traits are required (*c*_*2*_ > *c*_*1*_ > 0 and *b*_*1*_ > *b*_*2*_ > 0). In panels **(A)** and **(C)**, the cooperator for both traits (WT) and the cheater of the 1^st^ cooperative trait compete (*p*_*coop*_(0) = 0.9 and *p*_*ch1*_(0) = 0.1), while the cheater of the 2^nd^ cooperative trait is absent (*p*_*ch2*_(0)=0), whereas in panels **(B)** and **(D)** all three strains compete (*p*_*coop*_(0) = 0.8 and *p*_*ch1*_(0) = *p*_*ch2*_(0) = 0.1). The values that are given to the parameters of the simulations are: **(A)** *p*_*coop*_(0)= 0.9, *p*_*ch1*_(0)=0.1, *p*_*ch2*_(0)=0, *c*_*1*_=0.01, *b*_*1*_=3.4, *c*_*2*_=0, *b*_*2*_=0, *ω*_*0*_=0.2, *th*=0.8, *n*=30; **(B)** *p*_*coop*_(0)= 0.8, *p*_*ch1*_(0)=0.1, *p*_*ch2*_(0)=0.1, *c*_*1*_=0.01, *b*_*1*_=3.4, *c*_*2*_=0, *b*_*2*_=0, *ω*_*0*_=0.2, *th*=0.8, *n*=30; **(C)** *p*_*coop*_(0)= 0.9, *p*_*ch1*_(0)=0.1, *p*_*ch2*_(0)=0, *c*_*1*_=0.01, *b*_*1*_=1. 5, *c*_*2*_=0.025, *b*_*2*_=0.25, *ω*_*0*_=0.1, *th*=0.8, *n*=30; **(D)** *p*_*coop*_(0)= 0.8, *p*_*ch1*_(0)=0.1, *p*_*ch2*_(0)=0.1, *c*_*1*_=0.01, *b*_*1*_=1.5, *c*_*2*_=0.025, *b*_*2*_=0.25, *ω*_*0*_=0.1, *th*=0.8, *n*=30. Note that the values of parameters used in these simulations are chosen to reflect approximately the relation between the values observed in Figure 1, Figure 2. (For more detailed description please see Supplemental Information, Mathematical Model 2).

To test experimentally if quorum sensing regulation could indeed be the mechanism responsible for preventing fixation of *lasR* in the WT+*lasR* competitions, we repeated the propagation experiment shown in Figure 6A adding the quorum sensing autoinducer Acyl-homoserine lactone (AHL) N-3-oxododecanoyl-homoserine lactone (3OC_12_-HSL) to the culture medium. Addition of AHL abolishes the quorum sensing-dependent regulation of elastase, locking elastase production constitutively in the ON state. Remarkably, the addition of AHL allows the *lasR* mutant to expand throughout the competitions, and eventually reach fixation (Figure 6B), as the model without quorum sensing predicted (Figure 5A). Thus, regulation of the production of a public good by quorum sensing prevents full domination of the quorum sensing cheater, maintaining cooperation in populations. However, if the expanding cheater is affected in the production of a public good not regulated via quorum sensing (e.g. *pvdS*), this mutant can dominate the entire population.

In summary, the results obtained with our 3-way public goods model including quorum sensing (Figure 7) show that the dynamics observed in our propagation experiments (Figure 3) can be explained by the relationship between the cost of the different cooperative traits involved and a quorum threshold that regulates both costs and benefits of one of these traits. Figures S6E-I represent predictions, according to our model, for other possible scenarios with different relationships between the costs, which can be tested experimentally in the future (see Supplemental Information).

## Discussion

The classical experimental approach in sociomicrobiology has been to study one trait at a time. The simplicity of such an approach has allowed to substantially increase our understanding of the dynamics of cooperative interactions and revealed several mechanisms involved in the maintenance of cooperation [2,4,47]. In particular, the ability of *lasR* or *pvdS* mutants to behave as cheaters individually has been extensively documented [9,11,25–27,34,35,64,67,70–74], and these mutants are commonly isolated from bacterial populations colonizing CF lungs [41,45].

We established an experimental setup where WT cooperates in more than one trait: production of elastase via quorum sensing regulation and production of siderophore pyoverdine. Under conditions where the two traits are required, the *lasR* mutant can act a cheater for elastase but a cooperator for pyoverdine, whereas the *pvdS* mutant can cooperate for elastase production and cheats for pyoverdine production. Our results showed that, in this environment, the 3-way competitions result in a dominance of *pvdS* over both the WT and the *lasR* mutant. Presumably, this occurs because the advantage of the *pvdS* mutant (caused by not producing pyoverdine) is higher than that of *lasR* (for not producing elastase, and the other quorum sensing regulated goods) under conditions where the two traits are necessary, causing *pvdS* to be more fit than *lasR* in this environment (Figure 1C). As a consequence, the *pvdS* mutant can cheat on the *lasR* mutant (and on the WT) (Figure 2, Figure S1), dramatically affecting the outcome of the long-term competitions (Figure 3D).

The expansion of *pvdS* under conditions where the two traits are required prevents the drastic population collapse caused by invasion of *lasR* mutants observed in absence of *pvdS* (compare panels D and C of Figure 3). Even though the domination of *pvdS* mutant can also lead to a drop in the density of the population caused by the exhaustion of the public good, (Figure S3B and C), the decrease in cell density due *pvdS* domination is much less drastic than that of observed upon domination of *lasR* mutant (Figure 3A – C).

Interestingly, both *lasR* and *pvdS* mutants are stronger cheaters in the medium where either of their affected trait is required than in the medium where the two traits are necessary (Figure 2, panels A and D versus E and F). In the case of *lasR*, this difference is coherent with the lower growth yields reached under conditions where the two traits are required (Figure 1, A versus C), which allow fewer cell divisions and therefore milder cheating. The difference in cheating of *pvdS* cannot be ascribed to higher growth yields when only pyoverdine is required (Figure 1B and C). However, pyoverdine production per cell is significantly higher in the medium where only pyoverdine is required (Figure S2G-H), and this could possibly explain the boosting in the cheating by *pvdS* in this medium. A stronger iron depletion in the CAA + Transferrin medium is coherent with both the low iron content of the CAA mixture used in our experiments [75] and the iron-chelating capacity of casein [76,77], which could allow carryover of casein-bound iron to the medium and thus result in higher iron availability in the media with casein.

Our simple model assuming that the difference in costs and benefits of the cooperative traits involved proved to be sufficient to reasonably explain our experimental results. Therefore, the model allowed us to infer the general parameters governing social interactions beyond the particularities of the two mutants used in this study. Specifically, the mathematical model suggests that, in competitions among more than one social cheater under conditions where more than one trait is required (a scenario likely to be closer to the conditions in nature), the mutant for the trait with the highest cost is expected to dominate. Moreover, the degree of the decrease in population density caused by loss of cooperation due to exhaustion of the public good is determined by the benefit minus the cost difference of the trait affected. In case of a trait with high benefit-cost difference, a drastic collapse on the density of the population caused by the cheater in that trait is expected. In contrast, if the mutant for the trait with a low benefit-cost difference (as inferred for *pvdS*) dominates, a weak drop occurs. These scenarios that lead to different degrees of decrease in population densities could have very different consequences for the host in the context of infections.

Importantly, our results provide support for a dynamic view of cooperation and cheating that is dependent on both the genotypes present in the population and the environmental conditions. We demonstrated how changes in the abiotic environment can cause a social mutant to start or stop cheating or being cheated. Additionally, as shown here for the *lasR* mutant, quorum sensing regulation can also favor the maintenance of polymorphism, since such regulation alters the values of the cost and benefits of the traits as a function of the population density.

A better understanding of the interactions in polymorphic bacterial populations in complex environments not only helps to gain insights into key aspects of sociomicrobiology, but also can provide a theoretical framework for the development of new therapeutic strategies against bacterial populations where social mutants can invade [41,45]. In particular, our study provides relevant information about the biotic and abiotic conditions that favor the expansion of these mutants, which should be taken into account when considering strategies aiming to manipulate populations where this type of social interactions is taking place.

The potential effects of the appearance of *pvdS lasR* double mutants in settings similar to ours should also be considered since double mutants have the potential to occur *in vivo* [41]. Although we found no evidence for emergence of *pvdS lasR* double mutants within the period of the experiments reported here, in the course of longer propagations *pvdS lasR* double mutants generated by *de novo* mutations were identified (data not shown). The effects of these double mutants on the interactions described here should be investigated in the future. However, based on our results, we can speculate that double mutants, as full cheaters, should cause an accelerated collapse of the population.

A non-social explanation for the advantage of the *pvdS* mutant in triple co-cultures under conditions where the two traits are required was also considered given that, at least in *Pseudomonas fluorescens*, certain mutants defective in pyoverdine production have been reported to be better adapted even in environments where iron concentration is not low, and thus can be considered non-social mutations [78]. However, the fact that our *pvdS* mutant has a lower fitness than the WT in the low iron media and does not show any advantage in conditions where pyoverdine production is not necessary (Figure 1B and C, Figure 2B and H) rules out non-social adaptation as the reason for its advantage.

Collectively, our findings underline the need for including polymorphism in social phenotypes and multiple environmental conditions in experimental studies and mathematical models pertaining to cooperation in microbial populations. This need is further supported by recent theoretical and experimental studies showing that interactions between genetically and functionally interlinked cooperative traits can significantly affect the course of their social evolution [26,79]. Our results demonstrate that using experimental conditions that include more than one social trait can reveal complex and dynamic social roles in bacterial populations as well as their dependence on the environment. Understanding the dynamics of polymorphic populations in these complex environments provides insights into social interaction processes, expanding their relevance beyond sociomicrobiology, in addition to providing important knowledge for the development of novel therapeutic tools.

## Materials and Methods

### Bacterial strains

The strains used in this study were *Pseudomonas aeruginosa* WT strain PA01, PA01 *lasR* mutant harboring a gentamycin resistant gene inserted in *lasR (lasR*∷*GmR)* [80], and PA01 *pvdS* mutant harboring a gentamycin resistance gene resistance gene replacing the *pvdS* coding sequence (Δ*pvdS*∷*GmR*) [81]. For more detailed information, see Supplemental Information, supplementary methods section.

### Media and culture conditions

The medium where only elastase is required (iron-supplemented casein medium) contains casein (Sigma, Ref: C8654) (1% w/v) as the sole carbon and nitrogen source salts (1.18 g K_2_HPO_4_.3H_2_O and 0.25 g MgSO_4_.7H_2_O per liter of dH_2_O) and 50 μM of FeCl_3_. The medium where only pyoverdine production is required (iron-depleted CAA medium) contains the same salt solutions indicated above, low iron CAA (BD, Ref: 223050) (1% w/v) as the sole carbon source and 100 μg/ml of human apo-transferrin (Sigma, T2036) and 20 mM sodium bicarbonate to deplete available iron and induce pyoverdine production. The medium where both traits are needed (iron-depleted casein medium) is identical to the iron-supplied casein medium but instead of FeCl_3_ supplementation, this medium contains 100 μg/ml of human apo-transferrin (Sigma, T2036) and 20 mM sodium bicarbonate to deplete available iron and induce pyoverdine production. The medium where none of the traits is necessary (iron-supplemented CAA medium) contains the same salt solutions as the other media, low iron CAA (1% w/v) as the sole carbon source and 50 μM of FeCl_3_. All cultures were incubated in 15 ml falcons at 37°C with aeration (240 rpm, New Brunswick E25/E25R Shaker) for the incubation times indicated. Cell densities were estimated by measuring absorbance (Abs) at 600 nm (OD_600_) in a Thermo Spectronic Helios δ spectrophotometer.

### Determination of genotypic frequencies

Estimation of the frequencies of each strain in the co-cultures was performed by scoring fluorescence and colony morphology of colonies obtained from plating serial PBS dilutions of the cultures. For each individual sample, three aliquots (of 50μl - 200μl, as appropriated) were plated into LB agar plates, which were used as technical replicates. Then, CFU/ml were calculated by scoring different colony morphologies on each plate (with three technical replicate for each biological replicate). A stereoscope (Zeiss Stereo Lumar V12) with a CFP filter was used to distinguish pyoverdine producers, which are fluorescent, from the non-fluorescent *pvdS* mutants [72,82]. *lasR* mutant colonies have distinct colony morphology: smaller with smooth edges whereas elastase producers are larger with rugged edges [82]. To validate the phenotypic scoring all colonies used to determine the frequency from day 18 of the propagation experiments (Figure 3D) were tested by PCR with primers for the *lasR* and *pvdS* genes. The PCR data confirmed the phenotypic scoring with 100% accuracy.

### Measurement of relative mutant fitness

Relative fitness was used to determine the cheating capacity of each mutant as commonly used [72,83,84]. For both mutants (*lasR* and *pvdS*), the relative fitness of each mutant (*v*) was calculated as the change in frequency of the mutant over a period of 48 hours relative to the rest of the strains in the mixture, *i. e., v = fm*_*final*_.*fr*_*initial*_/*fm*_*initial*_.*fr*_*final*_ [72,83,84]. Where *fm* is the proportion of the mutant measured at the beginning of the competitions for *fm*_*initial*_, or after 48 hours of competition for *fm*_*final*_, and *fr* is the final proportions of the rest of the strains in the competitions at time = 0 (*fr*_*initial*_) or after 48 hours (*fr*_*final*_). As *fr* = (1 - *fm*), the relative fitness was determined using the following formula *v* = *fm*_*final*_ (1 - *fm*_*initial*_) / *fm*_*initial*_ (1 - *fm*_*final*_).

### Competition experiments

We propagated six replicates under four different conditions (Figure 3). Prior to start the competition experiments, all strains were inoculated, from frozen stocks, in medium containing 1% (w/v) casein and 1% (w/v) CAA in salts solution (same as in iron-supplied casein medium, described above) for 36 hours at 37°C temperature with shaking (240 rpm). Cells were then washed with PBS four times, to remove any residual extracellular factor. Next after measuring cell densities (OD_600_), cultures were normalized to OD_600_ = 1.0 and used to inoculate the various media as described in the text and figures. The different strains were diluted into fresh media, at different ratios as specified, to a starting initial OD_600_ = 0.05. For short term competitions the relative frequencies were determined by plating an aliquot of each culture at the beginning of the experiment (t = 0), and after 48 hours of incubation. For long-term competitions, the relative frequencies were determined at t = 0, and thereafter every 48 hours before each passage. At the end of every 48 hours 1.5 μl of each culture was transferred to 1.5ml of fresh medium (bottle-neck of 1/1000).

### Statistical analysis

Independent biological replicates were separately grown from the frozen stocks of each strain. Each figure (or figure panel) includes data from at least 6 biological replicates. The sample size (N), corresponds to the total numbers of independent biological replicates in each figure panel and is provided in the corresponding figure legends. The Mann-Whitney test which is a non-parametric test, was used because it does not account for normality and it is more suitable for the sample size used in each experiment (5<N<20). For multiple corrections, Kruskal-Wallis test with Dunn’s correction was used. For all statistical analyses we used GraphPad Prism 6 software (http://www.graphpad.com/scientific-software/prism).

## Acknowledgements

We thank Joana Amaro for technical assistance; João B. Xavier (Memorial Sloan Kettering), João Barroso-Batista, Rita Valente, and Jessica Thompson for suggestions and helpful comments on the manuscript, Jan Engelstädter (The University of Queensland, Australia) for his help while building the mathematical model, and Kevin Foster (University of Oxford, UK) for sending some of the strains used in this work. This work is supported by the Howard Hughes Medical Institute (International Early Career Scientist grant, HHMI 55007436). R.B. is supported by European Research Council (ERC-2010-StG_20091118) and by Fundação para a Ciência e Tecnologia (FCT) with a postdoctoral fellowship SFRH/BDP/109517/2015, Ö.Ö. is supported by Fundação Calouste Gulbenkian with a Doctoral Fellowship 01/BD/13, K.X. and I.G. are supported by FCT Investigator Program.

## Author Contributions

Conceptualization, Ö.Ö., K.X., I.G., and R.B.; Methodology, Ö.Ö., K.X., I.G., and R.B.; Investigation, Ö.Ö.; Writing - Original Draft, Ö.Ö. and R.B.; Writing – Review & Editing, Ö.Ö., K.X., I.G., and R.B.; Funding Acquisition, K.X. and I.G.; Resources, K.X. and I.G.; Supervision, K.X. and I.G.

